# SNHG16 promotes cell proliferation and inhibits cell apoptosis via regulation of the miR-1303p/STARD9 axis in renal cell carcinoma

**DOI:** 10.1101/2021.03.10.434814

**Authors:** Tao Cheng, Weibing Shuang, Dawen Ye, Wenzhi Zhang, Zhao Yang, Wenge Fang, Haibin Xu, Mingli Gu, Weiqiang Xu, Chao Guan

## Abstract

**Background:** Renal cell carcinoma (RCC) is a fatal malignant tumor with high morbidity. Numerous medical studies have suggested that long noncoding RNAs (lncRNAs) exert their biological function on various cancerous progresses. Herein, functions of lncRNA SNHG16 in RCC cells and the mechanism medicated by SNHG16 were investigated.

**Methods:** The expression levels of SNHG16 and its downstream genes in RCC cells and tissues were examined utilizing reverse transcription quantitative polymerase chain reaction analyses. Cell counting kit-8 and 5-Ethynyl-2’-deoxyuridine assays were carried out to evaluate the proliferation of RCC cells, and flow cytometry analyses were employed to determine the apoptosis of RCC cells. Western blot analysis was applied to examine protein levels associated with cell proliferation and apoptosis. The combination between SNHG16 and miRNA as well as miRNA and its target gene were explored by luciferase reporter, RNA pull down, and RNA immunoprecipitation assays.

**Results:** The significant upregulation of SNHG16 was observed in RCC tissues and cells. SNHG16 downregulation inhibited the proliferation and promoted the apoptosis of RCC cells. In addition, SNHG16 served as a competing endogenous RNA for miR-1301-3p, and STARD9 was a target gene of miR-1301-3p in RCC cells. SNHG16 upregulated STARD9 expression by binding with miR-1301-3p in RCC cells. Rescue assays validated that SNHG16 promoted RCC cell promotion and induced RCC cell apoptosis by upregulating STARD9 expression.

**Conclusions:** SNHG16 promotes RCC cell proliferation and suppresses RCC cell apoptosis via interaction with miR-1301-3p to upregulate STARD9 expression in RCC cells.

## Introduction

Renal cell carcinoma (RCC) is a prevalent malignant tumor that occurrs in genitourinary system with an increasing overall incidence [1]. The post-operative recurrence rate in RCC patients is up to 40% [2]. Most RCC patients preferred to choose traditional chemotherapy while the treatment effects may be limited due to the resistance of anticancer agents [3]. To enrich RCC treatments, the underlying molecular mechanisms in RCC are extensively explored.

Long noncoding RNAs (lncRNAs) are regarded as transcripts containing over 200 nucleotides and lacking the protein-coding capability [4]. Plenty of studies have validated that lncRNAs participate in multiple cellular activities, including cell proliferation, apoptosis, invasion, and migration in tumors [5-8]. Accumulating studies revealed that a multitude of abnormally expressed lncRNAs are involved in the development of RCC. For example, lncRNA LOC653786 promotes RCC cell growth and cell cycle by upregulating FOXM1 expression in RCC cells [9]. LncRNA ROR escalates the development of RCC via the miR-206/VEGF axis [10]. In addition, increasing evidence suggested that SNHG16 influences the development of various malignancies. For example, SNHG16 promotes the proliferation, migration and invasion of ovarian cancer cells via upregulation of MMP9 in ovarian cancer [11]. SNHG16 suppresses the proliferation of hepatocellular carcinoma cells by sponging miR-93 [12].

Mechanistically, numerous studies indicated that lncRNAs can be potential prognostic markers and interact with microRNAs (miRNAs) by functioning as competing endogenous RNAs (ceRNAs) in a variety of cancers [13-15]. To be specific, lncRNAs interacted with miRNAs to modulate the expression of target genes of miRNAs, thereby affecting malignant phenotypes of cancer cells, such as cell proliferation, migratory and invasive capacities, and apoptosis [16, 17]. As to RCC, LINC00461, SNHG4 and lncRNA HOTAIR were reported to function as ceRNAs to affect RCC development [18, 19]. In previous studies, SNHG16 was validated to function as a ceRNA for miR-140-5p to increase ZEB1 expression, thus facilitating cell proliferation, migration and EMT progress in esophageal squamous cell carcinoma [20]. SNHG16 contributes to the proliferative, migratory, and invasive abilities of pancreatic cancer cells by serving as a ceRNA for miR-218-5p to increase HMGB1 expression [21]. SNHG16 has prognostic values for patients with RCC, and its upregulation predicts unfavorable prognosis of RCC patients [22].

Herein, we hypothesized that SNHG16 was an oncogene in RCC and served as a ceRNA to regulate expression levels of downstream genes in RCC cells. SNHG16 expression in RCC tissues and cells, effects of SNHG16 expression on RCC cell proliferation and apoptosis, and the regulatory mechanism medicated by SNHG16 in RCC cells were under investigation. The study may provide novel insight into molecular studies of SNHG16 in RCC development.

## Materials and methods

### Bioinformatics analysis

The possible miRNAs binding to SNHG16 were predicted from starBase (http://starbase.sysu.edu.cn/) under the condition of “Degradome Data: low stringency (>=1)” and “Pan-Cancer: 6 cancer types” [23]. The possible target genes of miR-1301-3p were predicted from starBase under the condition of “CLIP Data: low stringency (>=1); Degradome Data: low stringency (>=1); Pan-Cancer: 10 cancer types; Program number: 1 program and predicted program: microT and miRmap”. STARD9 expression in kidney renal clear cell carcinoma (KIRC) tissues was also predicted from starBase.

### Clinical samples

Forty-five pairs of RCC tissues and noncancerous tissues were collected from RCC patients at The Second Affiliated Hospital of Bengbu Medical College. Before the surgery, patients had not received anticancer treatment, and written informed consent signed by patients had been obtained. All samples were immediately collected and reserved at −80°C after surgery. The Ethics Committees of The Second Affiliated Hospital of Bengbu Medical College approved this study.

### Cell culture

The normal human renal epithelial cell line HK2 and RCC cell line (A498, 786-O, Caki-1, and OSRC-2) were purchased from the American Type Culture Collection (ATCC; Manassas, VA, USA) and cultured in Roswell Park Memorial Institute (RPMI) 1640 medium (Thermo Fisher, Waltham, MA, USA) with 10% fetal bovine serum (Thermo Fisher). The cell culture was performed in a humidified atmosphere (37°C, 5% CO_2_).

### Cell transfection

Short hairpin RNAs against SNHG16 (sh-SNHG16#1/2) were transfected into RCC cells to knockdown SNHG16 expression, and sh-NC was set as a negative control. MiR-1301-3p expression was overexpressed utilizing miR-1301-3p mimics with NC mimics serving as a control. Complete sequence of STARD9 was inserted into the pcDNA3.1 vector to overexpress STARD9 expression. Lipofectamine 2000 (Invitrogen, Carlsbad, CA, USA) was applied for cell transfection in compliance with producer’s agreement. These synthetic plasmids and vectors were purchased from GenePharma (Shanghai, China). After two days, the efficiency of cell transfection was examined utilizing RT-qPCR.

### Reverse transcription quantitative polymerase chain reaction (RT-qPCR)

RNAs were extracted from A498 and 786-O cells using the TRIzol reagent (Takara, Otsu, Japan) and then reverse transcribed into cDNA using the TaqMan^™^ advanced miRNA cDNA synthesis kit (Thermo Fisher). SYBR Green PCR kit (Takara) was applied to conduct RT-qPCR. GAPDH is an internal control for SNHG16 and mRNAs while U6 is a reference for miRNAs. The results of RT-qPCR were analyzed by Step One Plus real-time PCR system (Applied Biosystems, Foster city, USA). The relative expression levels were estimated utilizing the 2^−ΔΔCt^ method. Primer sequences utilized in the study were listed below:

SNHG16

Forward: 5’-CTCTAGTAGCCACGGTGTG-3’

Reverse: 5’-GGGAGCTAACATTAAAGACATGG −3’

miR-1301-3p

Forward: 5’-TTGCAGCTGCCTGGGA-3’

Reverse: 5’-CTCTACAGCTATATTGCCAGCCAC −3’

STARD9

Forward: 5’-CCAGCTATTGAAGGAAGAGG −3’

Reverse: 5’-GAGAGGCTTCCATTAGAGC −3’

GAPDH

Forward: 5’-CCTCCTGTTCGACAGTCAG-3’

Reverse: 5’-CATACGA CTGCAAAGACCC-3’

U6 snRNA

Forward: 5’-TGCTATCACTTCAGCAGCA −3’

Reverse: 5’-GAGGTCATGCTAATCTTCTCTG −3’

### Western blot analysis

The protein contents were extracted utilizing an RIPA lysis buffer (Beyotime, Shanghai, China) and protease inhibitor (Roche, Basel, Switzerland). Next, the proteins were separated by SDS-PAGE for two hours and then were transferred to polyvinylidene fluoride membranes. Afterwards, the membranes were sealed with skim milk and incubated overnight with primary antibodies (Abcam, Cambridge, UK) at 4°C. Subsequently, secondary antibody (Abcam) was added to incubate these membranes for one hour at 37°C. GADPH was set as an internal control. At last, the protein bands were visualized using the enhanced chemiluminescent detection system (Thermo Fisher). The primary antibodies were anti-Cyclin A1 (ab53699; 1:500); anti-Cyclin B1 (ab32053; 1:3000); anti-PCNA (ab29; 1:1000); anti-CDK2 (ab101682; 1:100); anti-Bax (ab182733; 1:10000); anti-Bcl-2 (ab182858; 1:2000); and anti-GAPDH (ab8245; 1:1000).

### Cell counting kit 8 (CCK-8) assay

The transfected cells (1×10 ^3^ cells/well) were seeded to 96-well plates. After 12, 24, and 36 hours of incubation, each well was added with CCK-8 solutions (Dojindo, Kumamoto, Japan) for another 4 hours of incubation. Absorbance was observed by a EL340 microplate reader (BioTek Winooski, VT, USA) at 450 nm wavelength.

### 5-Ethynyl-2’-deoxyuridine (EdU) staining assay

The EdU Kit (RiboBio, Guangzhou, Guangdong, China) was utilized in this assay. The RCC cells (3000 cells/well) were added with 50 mM of EdU solution and fixed with 4% formaldehyde for 30 minutes. EdU reaction cocktail and DAPI (Beyotime, Nantong, China) were respectively used to incubate and counterstain the RCC cells. A fluorescence microscopy was employed to record the staining results. Images of five random fields were captured and the number of EdU-positive cells was counted.

### Flow cytometry analysis

Transfected cells were cultured in 6-well plates for 48 hours, and then washed and resuspended with phosphate buffer solution. After the cells were centrifugated at 300 g for 5 minutes, cells were stained with the Annexin V-FITC/PI apoptosis detecting kit (BD Biosciences, Franklin Lakes, NJ, USA) for 15 minutes. Finally, a flow cytometer (BD Biosciences) was employed to detect apoptotic rates.

### Subcellular fractionation assay

The nuclear and cytoplasmic parts of RCC cells were extracted utilizing a PARIS Kit (Thermo Fisher) based on instructions. RT-qPCR analyses were carried out to assess expression levels of SNHG16, GAPDH, and U6 in the cytoplasmic and nuclear components. GAPDH and U6 were controls for cytoplasmic and nuclear expression in cells.

### Luciferase reporter assay

The wild type (Wt) or mutant type (Mut) of SNHG16 sequence and the 3’-untranslated region (UTR) fragment of STARD9 containing miR-1301-3p binding site were subcloned into the pmirGLO vectors (Promega, Madison, WI, USA) to generate SNHG16-Wt/Mut vectors and STARD9-Wt/Mut vectors. MiR-1301-3p mimics or NC mimics were respectively cotransfected with SNHG16-Wt/Mut into A498 or 786-O cells with Lipofectamine 2000 (Invitrogen). MiR-1301-3p mimics, miR-1301-3p mimic + pcDNA3.1/SNHG16 or NC mimics were cotransfected with STARD9-Wt/Mut into A498 or 786-O cells. The luciferase activities were measured utilizing the luciferase reporter assay system kit (Promega).

### RNA immunoprecipitation (RIP) assay

The Magna RNA-binding protein immunoprecipitation kit (Merck Millipore, Billerica, MA, USA) was applied for the assay. A498 and 786-O cells were lysed using the RIP buffer. Next, the lysate was incubated with magnetic beads precoated with Ago2 antibody or Normal IgG. Afterwards, TRIzol reagent was utilized to extract the RNA and RT-qPCR analysis was performed to detect RNA expression.

### Statistical analysis

The data were shown as the means ± standard deviation (SD) and analyzed utilizing the SPSS 20.0 software (Chicago, IL, USA). Spearman correlation coefficient was utilized to analyze the correlation of gene expression. Student’s t test and one-way analysis of variance followed by Tukey’s *post hoc* test were performed to evaluate the variance of each group. All experiments were repeated thrice. Statistical significant was defined as p < 0.05.

## Results

### SNHG16 knockdown inhibits RCC cell proliferation and promotes cell apoptosis

RT-qPCR analysis revealed that SNHG16 expression in RCC tissues was higher than that in adjacent normal tissues (Figure 1A). Compared with SNHG16 expression in normal human renal epithelial cells, SNHG16 levels was markedly increased in RCC cells, especially in A498 and 786-O cells (Figure 1B). To investigate biological functions of SNHG16 in RCC cells, a range of loss-of-function experiments were conducted in A498 and 786-O with transfection of sh-SNHG16#1/2. Efficiency of SNHG16 knockdown was examined utilizing RT-qPCR analysis. After indicated treatment, a notable decrease of SNHG16 levels were observed in A498 and 786-O cells, indicating that SNHG16 expression was successfully knocked down (Figure 1C). Next, CCK-8 and EdU assays were carried out to detect the proliferation of RCC cells. SNHG16 knockdown contributed to the decrease in OD values and the number of EdU positive cells, showing that the proliferation of RCC cells was suppressed (Figure 1D-1E). Flow cytometry analysis revealed that SNHG16 downregulation induced the apoptosis of A498 and 786-O cells (Figure 1F). According to western blotting, the protein levels of Cyclin A1, Cyclin B1, PCNA, CDK2 and Bcl-2 were significantly decreased while the Bax protein expression was markedly upregulated in RCC cells with transfection of sh-SNHG16#1/2 (Figure 1G).

**Figure 1.**
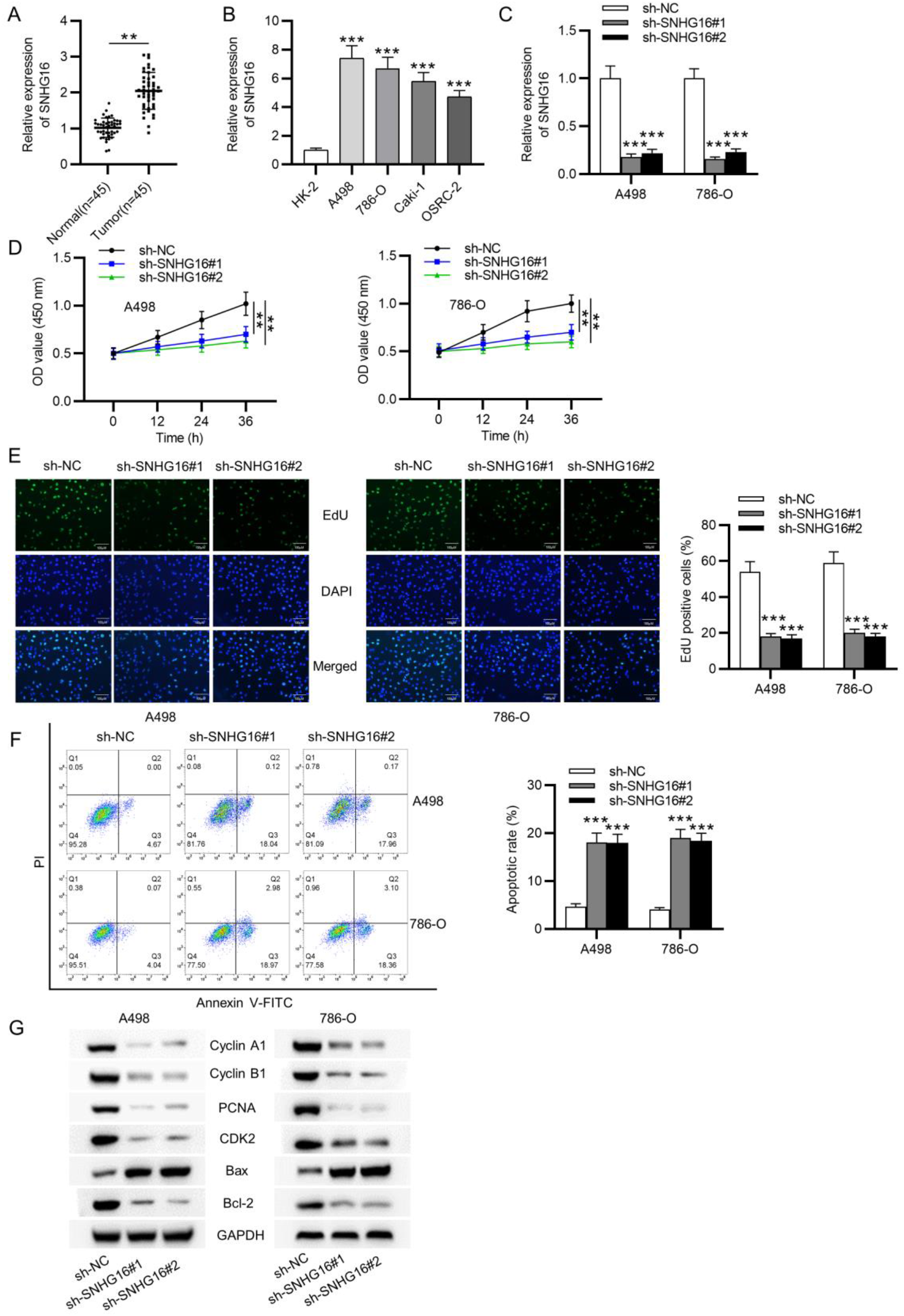
SNHG16 knockdown inhibits RCC cell proliferation and promotes cell apoptosis. (A) RT-qPCR analysis was employed to assess SNHG16 expression in RCC tissues and normal renal tissues (n=45). (B) SNHG16 expression levels in normal renal cells and RCC cells were measured by RT-qPCR analysis. (C) Efficiency of SNHG16 knockdown in A498 and 786-O cells was determined by RT-qPCR analysis. (D-E) CCK-8 and EdU assays were employed to probe the proliferation of RCC cells with transfection. (F) Flow cytometry analysis was performed to probe apoptotic rate of RCC cells after SNHG16 expression being knocked down. (G) Effects of SNHG16 silencing on protein levels of Cyclin A1, Cyclin B1, PCNA, CDK2, Bax and Bd-2 were probed utilizing western blot analysis. ** p < 0.01, *** p < 0.001.

### SNHG16 interacts with miR-1301-3p in RCC cells

To explore the localization of SNHG16 in RCC cells, subcellular fractionation assay was conducted, showing that SNHG16 was primarily distributed in the cytoplasm of RCC cells (Figure 2A). Next, startBase website was employed to predict potential miRNAs binding to SNHG16, and five candidate miRNAs (miR-196b-5p, miR-140-5p, miR-193a-5p, miR-1301-3p, miR-3690) were identified (Figure 2B). Among these candidate miRNAs, miR-1301-3p expression was downregulated in all the RCC cells (Figure 2C). Next, miR-1301-3p expression in RCC tissues was probed utilizing RT-qPCR analysis, revealing a significant decrease of miR-1301-3p expression in tumor tissues (Figure 2D). Spearman correlation coefficient was probed to identify the expression correlation between SNHG16 expression and miR-1301-3p expression. We found that miR-1301-3p expression was negatively correlated to SNHG16 expression in RCC tissues (Figure 2E). Moreover, SNHG16 knockdown significantly upregulated miR-1301-3p expression in A498 and 786-O cells, as shown in RT-qPCR (Figure 2F). Subsequently, we explored the combination between SNHG16 and miR-1301-3p. The possible binding site between SNHG16 and miR-1301-3p was predicted from starBase (Figure 2G). RT-qPCR examined efficiency of miR-1301-3p overexpression in A498 and 786-O cells (Figure 2H). A luciferase reporter assay suggested that the luciferase activity of miR-1301-3p-Wt was remarkably increased by SNHG16 knockdown while no significant changes in luciferase activity were detected in miR-1301-3p-Mut vectors (Figure 2I). According to an RIP assay, both SNHG16 and miR-1301-3p expression were increased in Ago2 antibody groups compared with these in IgG control groups (Figure 2J).

**Figure 2.**
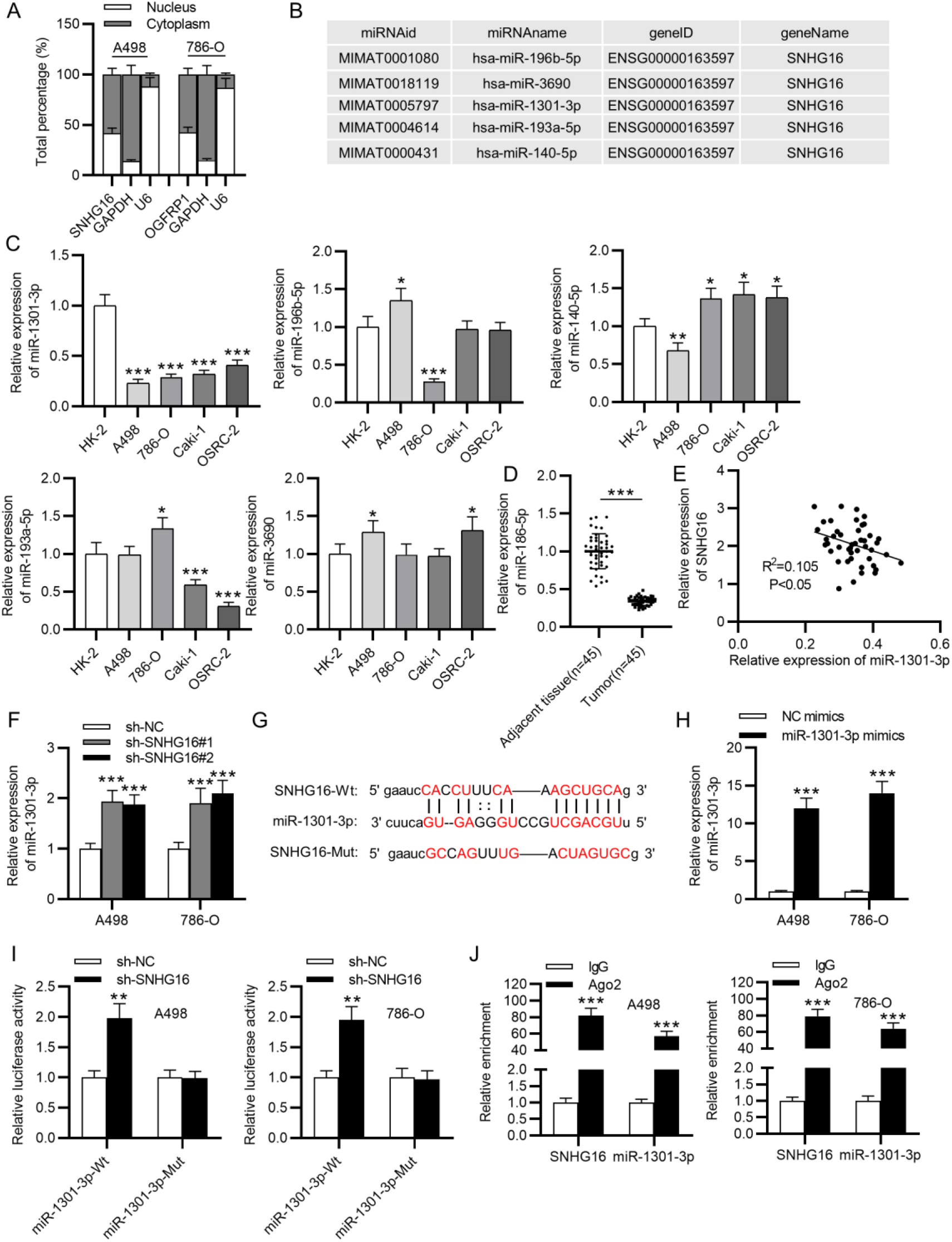
SNHG16 interacts with miR-1301-3p in RCC cells. (A) The subcellular fractionation assay was conducted to detect SNHG16 distribution in RCC cells. (B) MiRNAs combining with SNHG16 were predicted from the starBase website. (C) RT-qPCR analysis was performed to assess candidate miRNA expression in RCC cells and normal renal cells. (D) MiR-1301-3p expression in RCC tissues and normal nontumor tissues were examined by RT-qPCR analysis. (E) The correlation between SNHG16 expression and miR-1301-3p expression in RCC tissues were identified utilizing Spearman correlation coefficient. (F) Effects of SNHG16 expression on miR-1301-3p expression in RCC cells were assessed by RT-qPCR analysis. (G) The putative binding site between SNHG16 and miR-1301-3p was predicted from starBase. (H) Efficiency of miR-1301-3p overexpression was evaluated by RT-qPCR analysis. (I-J) Luciferase reporter and RIP assays were conducted to explore the interaction between SNHG16 and miR-1301-3p in RCC cells. * p < 0.05, ** p < 0.01, *** p < 0.001.

### STARD9 is a downstream target gene of miR-1301-3p

Possible target genes of miR-1301-3p were predicted utilizing starBase. Six candidate genes (GABARAPL1, STARD9, MYH11, MYH9, EDEM1 and CLU) were identified (Figure 3A). Compared with expression levels of other candidate genes, STARD9 expression was significantly downregulated due to miR-1301-3p overexpression in A498 and 786-O cells (Figure 3B). The high expression of STARD9 in KIRC tissues was predicted from starBase website (Figure 3C). The starBase website also provided possible binding site between miR-1301-3p and STARD9 (Figure 3D). According to a luciferase reporter assay, the luciferase activity of STARD9-Wt was markedly decreased due to miR-1301-3p overexpression, and the decrease in luciferase activity was reversed by SNHG16 overexpression. In addition, no significant changes in luciferase activity were examined in STARD9-Mut groups (Figure 3E). The findings suggested that STARD9 bound with miR-1301-3p at the predicted site. Compared with the expression in normal control IgG group, expression levels of SNHG16, miR-1302-3p and STARD9 were all significantly enriched in the Ago2 antibody group, as shown in RIP assays (Figure 3F). The results suggested that SNHG16, miR-1302-3p and STARD9 were enriched in the RNA-induced silencing complexes. Moreover, the mRNA expression of STARD9 was reduced by SNHG16 knockdown according to RT-qPCR analysis (Figure 3G). Western blot analysis implied that SNHG16 knockdown or miR-1301-3p overexpression downregulated the protein levels of STARD9 (Figure 3H).

**Figure 3.**
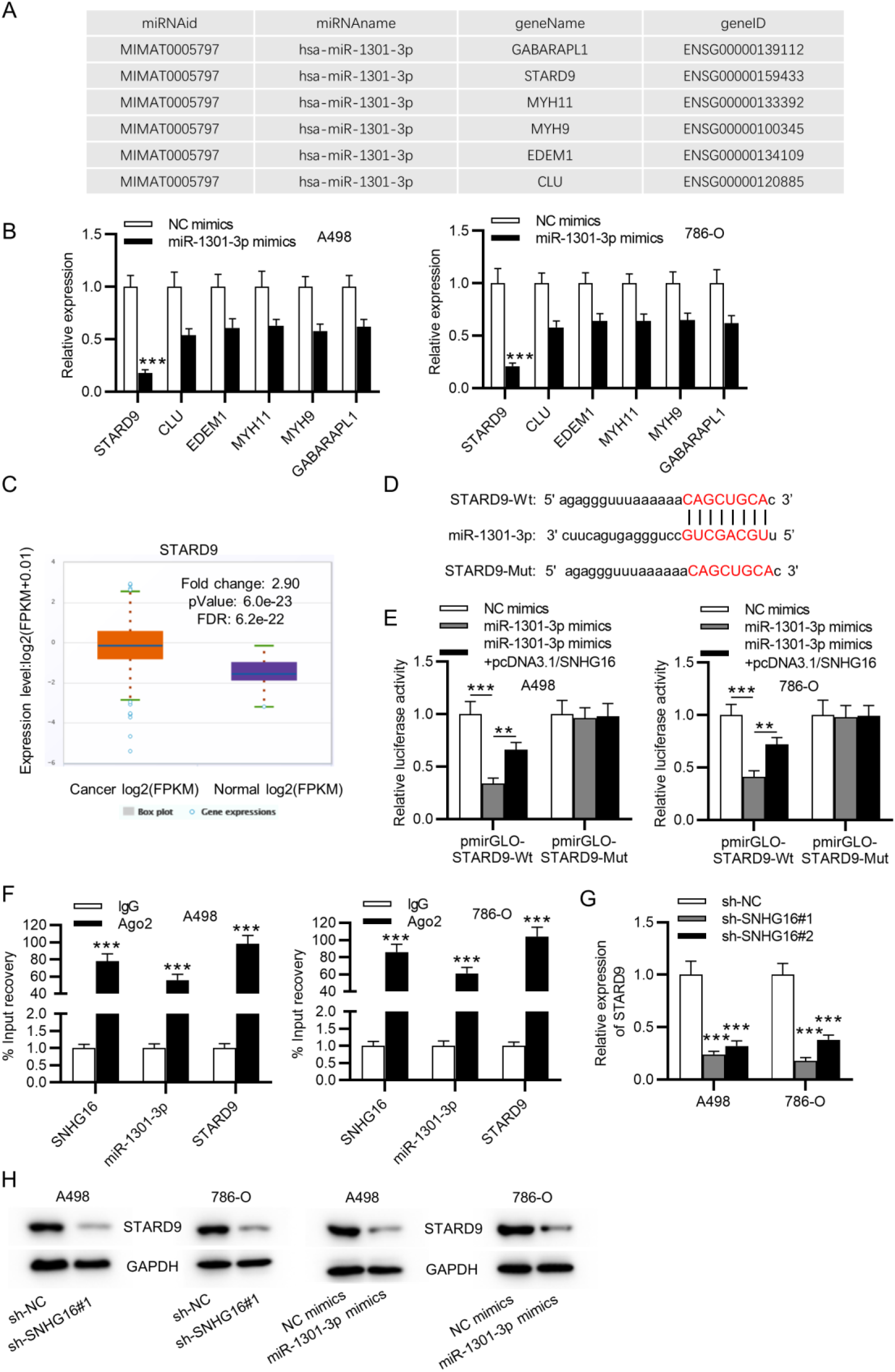
STARD9 is a target gene of miR-1301-3p. (A) The starBase website provided potential mRNAs that interact with miR-1301-3p. (B) STARD9 expression was significantly downregulated by miR-1301-3p overexpression compared with expression levels of other genes. (C) The starBase website predicted that STARD9 expression was significantly upregulated in KIRC tissues. (D) The possible binding site between miR-1301-3p and STARD9 was predicted from starBase. (E) The combination between miR-1301-3p and STARD9 was measured by the luciferase reporter assay. (F) The relationship among SNHG16, miR-1301-3p and STARD9 was further explored by the RIP assay. (G) Effects of SNHG16 expression on mRNA expression of STARD9 were examined using RT-qPCR analysis. (H) Western blot analysis was employed to assess influences of SNHG16 and miR-1301-3p expression on protein levels of STARD9 in cell s. ** p < 0.01, *** p < 0.001.

### STARD9 exhibits high expression in RCC cells and tissues

RT-qPCR analysis revealed that STARD9 expression in the RCC cells was upregulated compared with that in normal renal cells (Figure 4A). Compared with STARD9 expression in adjacent nontumor tissues, STARD9 expression in tumor tissues was significantly upregulated (Figure 4B). Based on Spearman correlation coefficient, SNHG16 expression was positively correlated with STARD9 expression while miR-1301-3p was negatively correlated with STARD9 expression in RCC tissues (Figure 4C-4D).

**Figure 4.**
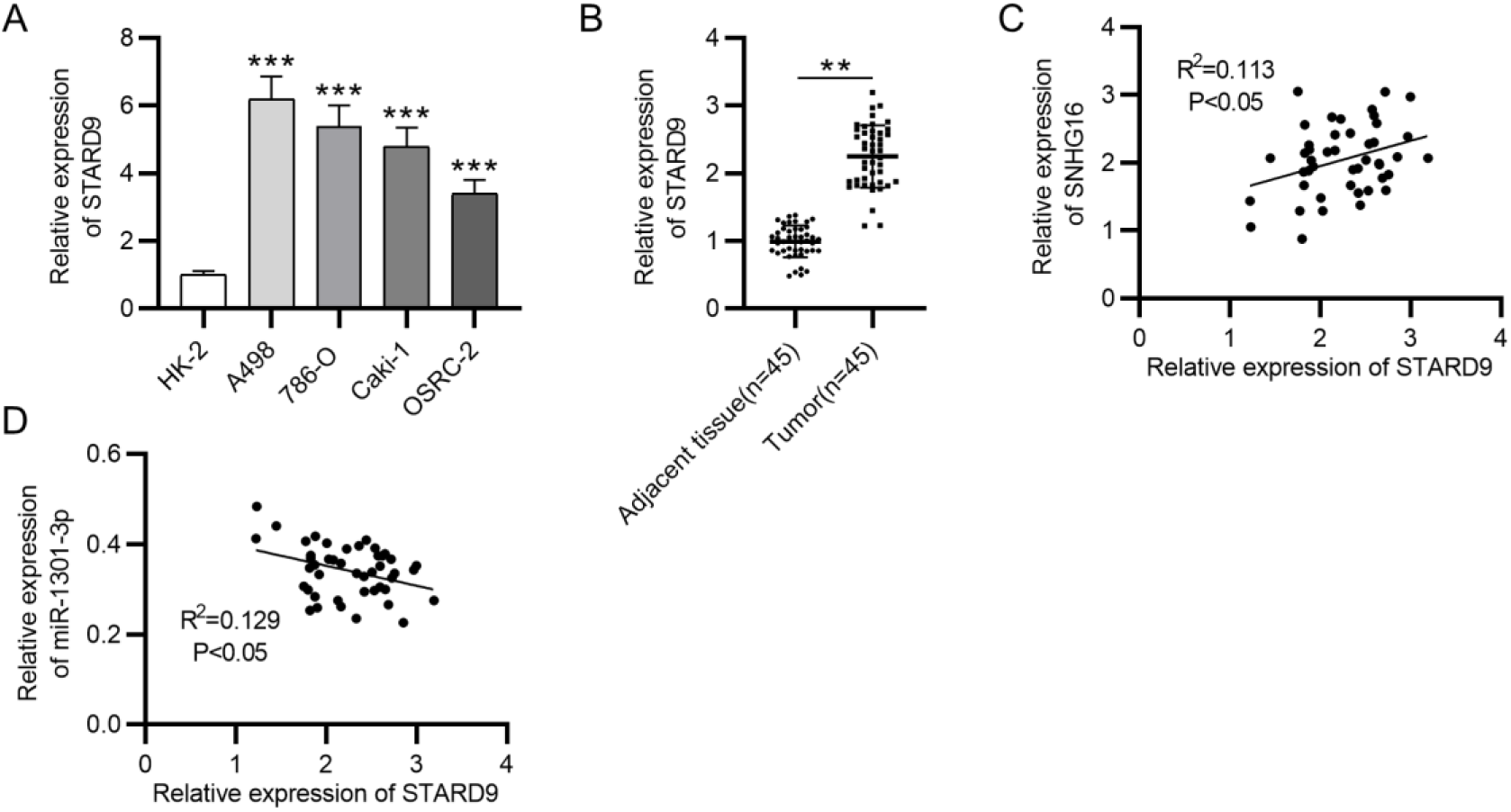
STARD9 exhibits high expression in RCC cells and tissues. (A) STARD9 expression levels in RCC cells and normal renal cells were probed by RT-qPCR. (B) STARD9 expression was relatively increased in RCC tissues according to RT-qPCR. (C-D) The correlation between STARD9 expression and SNHG16 expression or miR-1301-3p expression in RCC tissues was identified utilizing Spearman correlation coefficient. ** p < 0.01, *** p < 0.001.

### STARD9 overexpression reverses effects on RCC cell proliferation and apoptosis mediated by SNHG16 depletion

Whether SNHG16 promotes cell proliferation and promotes cell apoptosis by upregulation of STARD9 expression in RCC cells were verified by rescue assays. STARD9 levels were overexpressed in A498 and 786-O cells with transfection of pcDNA3.1/STARD9 (Figure 5A). STARD9 overexpression rescued suppressive effects on RCC cell proliferation mediated by SNHG16 silencing as shown in CCK-8 and EdU assays (Figure 5B-5D). A flow cytometry assay revealed that STARD9 overexpression reversed the enhancement of cell apoptosis caused by SNHG16 knockdown (Figure 5E). Compared with the expression in control group, the protein levels of Cyclin A1, Cyclin B1, PCNA, CDK2, and Bcl-2 were decreased while Bax protein levels were upregulated after SNHG16 depletion, and STARD9 overexpression reversed the changes in protein levels in RCC cells (Figure 5F).

**Figure 5.**
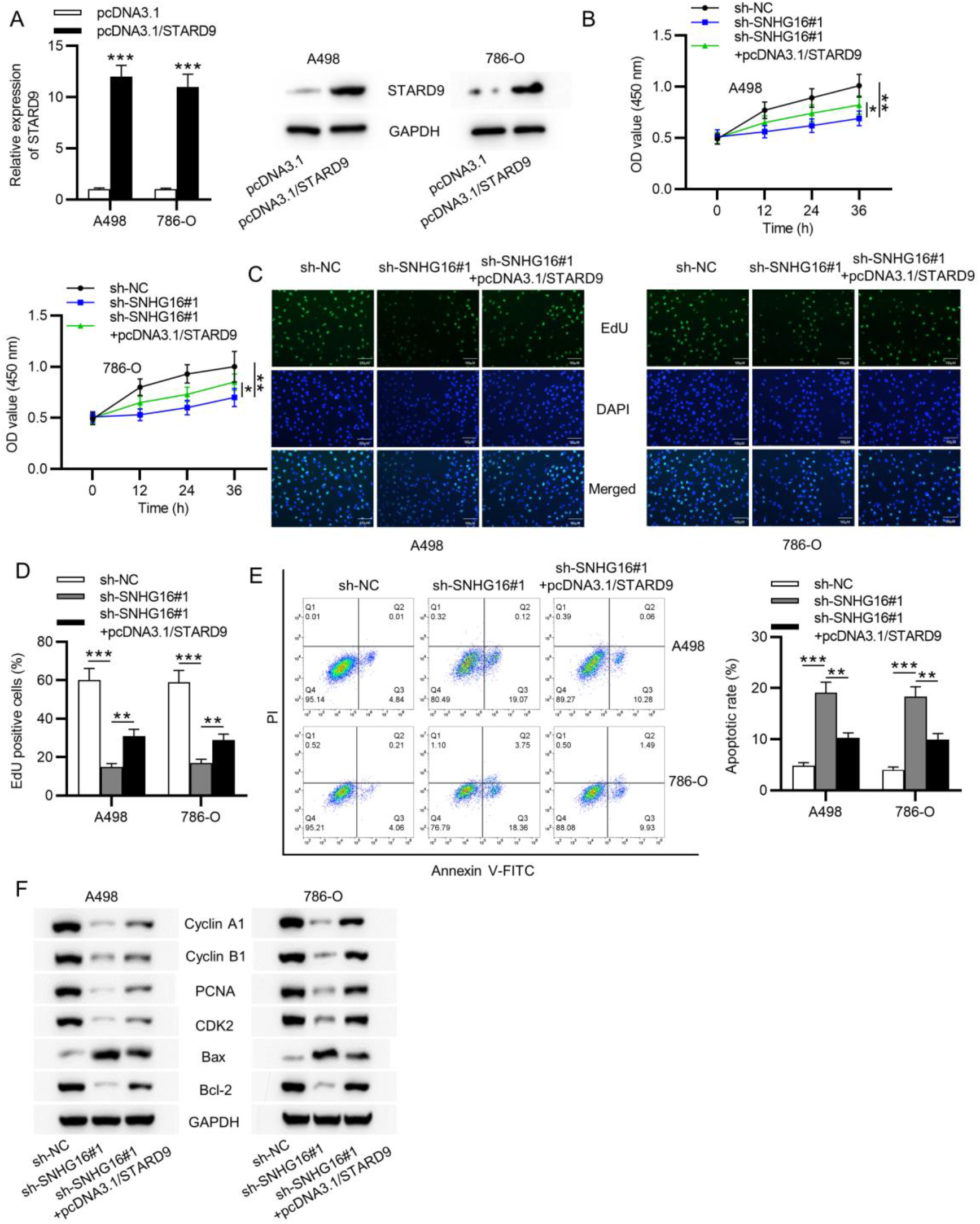
STARD9 overexpression reverses effects on RCC cell proliferation and apoptosis mediated by SNHG16 depletion. (A) RT-qPCR was applied to measure the efficiency of STARD9 overexpression in RCC cells with transfection of pcDNA3.1/ STARD9. (B-D) Effects of SNHG16 knockdown and STARD9 overexpression on the proliferation of RCC cells were explored utilizing CCK-8 and EdU assays. (E) The flow cytometry analysis was applied for detecting apoptotic rate of RCC cells with transfection. (F) Western blot analysis was employed to evaluate the protein levels of key factors related to cell proliferation and apoptosis. * p < 0.05, ** p < 0.01, *** p < 0.001.

## Discussion

SNHG16 was previously reported to be oncogenic in ovarian cancer [11], esophageal squamous cell carcinoma [20] and pancreatic cancer [21] while serve as an antioncogene in hepatocellular carcinoma [12]. In our study, SNHG16 was significantly upregulated in RCC cells and tissues, and was primarily located in the cytoplasm of RCC cells. SNHG16 knockdown inhibited cell proliferation and facilitated cell apoptosis. Additionally, SNHG16 served as a ceRNA against miR-1301-3p in RCC cells.

MiRNAs are short noncoding RNAs containing 20-24 nucleotides, which have been proved to play the oncogene role, function as tumor suppressor or regulate gene expression in numerous cancers [24]. MiR-1301-3p has been verified to take part in the progression of some cancers. For example, miR-1301-3p inhibits the proliferation of breast cancer cells by targeting ICT1 in human breast cancer [25]. MiR-1301-3p promotes cell expansion by suppressing GSK3β and SFRP1 expression in prostate cancer [26]. In addition, miR-1301-3p has been verified to be implicated in the ceRNA mechanism in different cancers. LncRNA ABHD11-AS1 serves as a ceRNA for miR-1301-3p to increase STAT3 expression in papillary thyroid carcinoma [27]. LncRNA MIAT facilitates cell migration and invasion in esophageal squamous cell carcinoma by acting as a ceRNA for miR-1301-3p to increase INCENP expression [28]. In this study, miR-1301-3p expression was significantly decreased in RCC cells and tissues. SNHG16 expression was negatively correlated with miR-1301-3p expression in RCC tissues. SNHG16 served as a ceRNA to upregulate STARD9 expression via interaction with miR-1301-3p in RCC cells.

Steroidogenic acute regulatory protein–related lipid transfer domain containing 9 (STARD9) is a member of the Kinesin-3 family, and its absence may cause apoptotic cell death, mitotic arreast and unabling to congress chromosomes [29]. Since STARD9 depletion exerts effects on the division of diverse cancer cells, such as colorectal carcinoma cells, non-small cell lung carcinoma cells, melanoma cells and cervical adenocarcinoma cells, STARD9 is believed to become a candidate target for enriching cancer therapeutics [29]. Based on bioinformatics analysis, STARD9 expression was predicted to be upregulated in KIRC tissues. In our study, STARD9 showed high expression in RCC tissues and cells. According to Spearman correlation coefficient, STARD9 expression was negatively correlated with miR-1301-3p expression while positively correlated with SNHG16 expression in RCC tissues. STARD9 was verified to be a target gene of miR-1301-3p, and miR-1301-3p directly interacted with the 3’UTR of STARD9 in RCC cells. Under the ceRNA mechanism, mRNAs can reverse the miRNAs-induced suppressive effects on them [13]. In our study, STARD9 overexpression reversed the suppressive effects on RCC cell proliferation as well as the promotion of RCC cell apoptosis induced by SNHG16 depletion.

In conclusion, SNHG16 promotes RCC cell proliferation and induces RCC cell apoptosis by acting as a ceRNA for miR-1301-3p to upregulate STARD9 expression. Our study may provide new insight into the regulatory mechanism of SNHG16 at the post-transcriptional level in RCC development.

## Acknowledgement

We appreciate all participants for their contribution to the study.

## Conflicts of interest

None.

